# CYTOKINE-BASED MULTI-EPITOPE PROTEIN FOR BINDING TO CCR7-POSITIVE CELLS

**DOI:** 10.1101/2022.11.22.517544

**Authors:** Maria Beihaghi, Hasan Marashi, Reihaneh Karimi, Mohammad Reza Beihaghi, Masoud Chaboksavar, Mahsa Zabetian

**Affiliations:** College of Agriculture, Ferdowsi University of Mashhad, Mashhad, Iran; Department of Biology, Kavian Institute of Higher Education, Mashhad, Iran; School of Science and Technology, The University of Georgia, Tbilisi, Georgia; Department of Pharmacology,Institute of Neurocience and Physiology, University of Gothenburg, Gothenburg, Sweden; Department of Psychology, Sheffield Hallam University, Sheffield, United of Kingdom

**Keywords:** Multiepitope protein, CCR7 receptor, Recombinant protein, Cytokine genes

## Abstract

Drugs based on cytokine genes increase the body’s immunity against cancer and several viral diseases. Cytokines such as CCL21 and CCL19 specifically bind to the CCR7 receptor and have anti-tumor properties and a prognosis of tumorigenesis. An epitope of IL-Iβ is also involved in various cellular activities such as neutrophil activation, T and B lymphocyte cell production, antibody production, and fibroblast proliferation. They bind to secretory proteins without any inflammatory response. GM-CSF adjuvant is one of the growth factors of white blood cells (WBC) that stimulates stem cells to produce granulocytes and myocytes. In our project, we designed and synthesized codon-optimized multiepitope genes construct from human genes. The molecular dynamics (MD) simulation and binding affinity of this recombinant protein and CCR7 receptor were examined through in silico analyses. This construct was introduced into the pET-28a vector that was cloned in Ecoli to explore this recombinant protein. The purified multiepitope protein produced a strong signal in Dot-blot, SDS-PAGE, and Western-blot assays comparable to the positive control. The assessments of FTIR measurement and MALDI-TOF MS displayed that synthetic gene constructs correctly be expressed in E. coli. We also investigated the potential activity of the purified multiepitope protein in stimulating migration and proliferation of MCF7CCR7+ cancer cell line using wound healing assay. Also, in this study, the MTT method was used to determine the half-maximum inhibitory concentration (IC50) of all multiepitope protein concentrations. We used Agarose assay on PBMCsCCR7+ to observe whether PBMC cells and chemoattractants attract to each other. Also, we found that intraperitoneal injection of recombinant antigen affected the level of WBC in BALB/c mice, and the level of WBC in tumor mice increased significantly compared to healthy mice. Our project aims to produce the first multi-epitope vaccine with many beneficial advantages such as low-cost price and any major or significant complications that can be used as biomarkers for cancer screening and prognosis tests to immunize the patient before chemotherapy.

## 1. Introduction

Today we are witnessing the spread of cancer so different methods have been used so far to treat cancer diseases(1); Chemotherapy is one of the methods used especially to cure cancer, which unfortunately causes some negative side effects such as nausea, weakness and hair loss (1, 2). Side effects are problems that arise in humans through treatment. These complications occur because the goal of cancer treatment is to destroy cancer cells, and in this fight, some healthy cells are also damaged(3). For example, Herceptin is one of the drugs prescribed to most cancer patients (4, 5), causes many side effects, and weakens the immune system in addition to its high cost, (4, 6). As a result, utilizing a compound that can strengthen and improve the immune system without any side effects before cancer treatment like chemotherapy is significant (7, 8). Currently, there are different animal-derived drugs as immunotherapy agents based on CCL21, CCL19, GM-CSF, and CCR7 receptors to strengthen the patient’s immune system by stimulating the immune system(9, 10). Specifically, the CCR7 and its ligands play an essential role in transporting T-lymphocytes and antigen-presenting cells, such as dendritic cells (DCs) to the lymph nodes. As a result, CCL19 and CCL21, together with the CCR7 receptor, are generally essential in strengthening the immune system and self-balance in physiological and pathophysiological situations(11, 12). Also, CCL21 now recognizes as an effective drug in the treatment of AIDS and cancer in international cancer institutes (13). However, these immunotherapy agents have several limitations. For example, CCL21 only affects on CTLs and T-lymphocytes and reduces regulatory T-cell expression, but it has no role in activating macrophages, B-lymphocytes and neutrophils(14). While the CCR7 receptor is present on surfaces of a wide range of immune cells, including T lymphocytes, B lymphocytes, neutrophils, and macrophages and it can be a suitable target for activating immune systems(15, 16). Therefore, in this project, to treat and prevent cancer and viral diseases, the production of recombinant multi-epitope drugs has been designed and produced. In addition to the CCL21 sequence, it contains fragments of IL1β, CCL19, and GMCSF amino acid sequences(17). By placing epitopes related to CTLs and B lymphocytes in the desired structure to activate B and T lymphocytes(18), neutrophils and macrophages simultaneously to use as a potent adjuvant for the treatment of breast cancer, especially in combination with *Her2 / neu* gene expression in inducing Th1 immune response, which affects most drugs improve the disease and increase immunity(16, 19). This multi-epitope binds specifically to the CCR7 chemokine receptor and has antitumor properties that can be used to predict tumor genesis in cancer patients. Increased CD8+T cells detect and reduce the progression of viral diseases such as HIV and COVID-19. The high expression of these genes acts as biomarkers in various diseases such as atherosclerosis, various cancers, bone disorders, and other inflammatory factors such as asthma, and viral infections such as HIV, and pneumonia(18, 20). Therefore, this recombinant protein can be produced at a very low cost, without any special side effects, to strengthen and improve the immune system to immunize the patient before chemotherapy (21, 22). So far, many immunotherapy drugs have been produced for this purpose that can strengthen the immune system of the patient by stimulating the immune system. We hope that if we succeed in the clinical phase while saving foreign exchange costs for the import of expensive drugs, the commercial production of this drug will also achieve the goal of monetization. Also, there is a need for producing recombinant cytokine-based multi-epitope drugs that can simultaneously stimulate a patient’s innate and adaptive immune systems without causing an autoimmune reaction. Furthermore, there is a need for a helpful biomarker for cancer screening and prognosis tests(23, 24). So, this project aims to produce the first multi-epitope vaccine with four epitopes such as CCL21, CCL19, IL-Iβ and GM-CSF for simultaneous activation of T and B cell lymphocytes, macrophages and neutrophils required to transfer protein to the target tissue. Additionally, we want to produce specific sites and linkers to stabilize and activate proteins. We hypothesize that transgenic bacteria-derived antibodies can be a viable alternative to other conventional antibodies(25). So far there have been no reports of production of cytokine gene-based multi-epitope drugs that can simultaneously stimulate a patient’s immune cells without causing an autoimmune reaction, and no recombinant drug has been developed. Therefore, there is a need to produce a multi-epitope drug based on cytokine genes with high immunogenicity and specificity to destroy cancer cells.

## 2. Material and Method

### 2.1 Evaluation of recombinant multi-epitope by silico tools

#### 2.1.1 Engineering and designing gene structures

To produce and manufacture the multi-epitope vaccine and to increase the ability of the vaccine to bind to the CCR7 receptor, a gene construct was designed and manufactured artificially. To make the vaccine, the amino acid sequences of CCL21, CCL19, IL-Iβ, and GMCSF were taken from NCBI database and were identified by the relevant BCpreds (http://ailab-projects1.ist.psu.edu:8080/bcpred/),BepiPred2.0(https://services.healthtech.dtu.dk/service.php?BepiPred2.0),ABCpreds(http://crdd.osdd.net/raghava/abcpred/),SVMTrip(http://sysbio.unl.edu/SVMTriP/), and MAPPP online servers,(http://mendel.stanford.edu/SidowLab/downloads/MAPP/index.html), B-cell and T-cell epitopes(15, 17). Peptide signal regions were selected using Signal P software and deleted to shorten the sequence of the gene construct. In order to achieve high expression and find a suitable region for ribosome binding, the expression enhancing sequence (Kozak sequence) was added to the 5’ end, and then the Histidine tag sequence (6xHis) was added to the 3’ end of the gene. The purpose of adding a polyhistidine tag is to help to purify the protein in the final stages. Therefore, in the gene construct, the GM-CSF was connected to the CCL19 through a helical linker (EAAAK), a beta-defensins that reduces interaction with other recombinant protein domains. The CCL19 was connected to the CCL21 through a furine protease-sensitive linker (RRVR). Also, the CCL21 was connected to the truncated IL-1β through a cathepsin B-sensitive linker (GPGPG). The IL-1β was directly connected to the rat KC chemokine without a linker. Moreover, the rat KC chemokine was connected to the polyhistidine tag (6x His tag) through an HIV protease-sensitive linker (RVLAEA). After designing the exemplary cytokine-based multi-epitope protein, its physicochemical characteristics were determined using in-silico tools(26).

#### 2.1.2. Molecular modeling and Molecular dynamic simulation of cytokine-based multi-epitope protein

Initially, due to the fact that the three-dimensional structure of the recombinant protein is not available, for this reason, the comparative modeling method was used to create the three-dimensional structure of the recombinant protein. The corresponding recombinant protein is composed of several proteins including GM-CSF, CCL21, CCL19 and IL1β, and for this reason, the mentioned proteins were used to model this recombinant protein (if there is a three-dimensional structure in the PDB database). Comparative modeling is one of the best ways to obtain the three-dimensional structure of the target protein. In this method of modeling, three-dimensional structures whose sequences are very similar to the target sequence are used as a model. One of the best software used for comparative modeling is MODELLER software. In this section, modeller 9.24 software was used for 3D modeling of the target wax. Also to find changes in the orientation of different amino acids over time, molecular dynamics simulations was used (27). In this system, each atom is dynamic, which causes movement in the entire protein structure, which can optimize the 3D structure of the recombinant multiepitope modeling protein.(28, 29). Motion can eliminate a series of interactions in the structure of proteins and thus the formation of new interactions, which can optimize the structure of the proteins, so a change in motion causes a change in structure and thus a change in protein function.(30). The MD simulation was performed by using Gromacs 2019.6 software. Input target structures were prepared with ff99SB force field. The disulfide bonds and correct hydrogen status of histidine amino acids was defined for all proteins. The surface charge of the structure was neutralized by adding chlorine ions(30). The target protein was placed in a layer of 8-angstrom thick TIP3P water molecules inside an octahedron box using gmx solvate software. Molecular dynamics simulations were performed at 37 °C for 100 nanoseconds. The SHAKE algorithm was used to increase computational speed and to limit the bonds involved in the hydrogen atoms. Finally, some analyses were used to evaluate the stability structure of target protein during the simulation. One of the best analyses is that the root means square deviation (RMSD) and Radius of Gyration change over time (30). The RMSD squares between the structures created during the molecular dynamics simulation in the time dimension is a suitable and common standard to ensure the structural stability of the protein. Therefore, the RMSD changes related to alpha carbon atoms of the protein during the simulation time (100 ns) relative to the original structure were calculated and extracted. Radius of Gyration is one of the important parameters in the study of changes in protein size during the simulation of molecular dynamics.

#### 2.1.3. Molecular Docking and simulation of recombinant protein with CCR7 receptor

In this step, HADDOCK software was used to investigate the interaction of the recombinant protein with CCR7 protein. In this software, biochemical and biophysical information obtained from laboratory methods, etc. are used to predict the manner of interaction(31). Using molecular dynamics simulation technique, this program performs protein-protein docking in a completely flexible way(32). To limit the volume of docking calculations, residues that are directly involved in the connection are identified. In laboratory methods, methods such as targeted mutagenesis, chemical shift perturbation, etc. are used to obtain the amino acids involved in the interaction(33). In this method, amino acids in which the level of water exposure is more than 50% are considered as active amino acids. Then, according to the mentioned protocol, recombinant active amino acids of protein were identified(34). Also, extracellular amino acids of CCR7 protein were considered as active amino acids. After performing molecular docking, the best selected complex was used as an input to simulate molecular dynamics. The protocol mentioned in Section 5 was used to place the complex inside the membrane and perform molecular dynamics simulations. The only difference is the number of phospholipids, water molecules and the number of sodium and chlorine ions used. In this part, the number of POPC phospholipids is equal to 179 molecules, the number of aqueous molecules is 20443 and the numbers of sodium and chlorine atoms are 54 and 89 atoms, respectively. Also, the simulation time is 100 nanoseconds. In this section, the RMSD parameter was used to study the stability of the recombinant protein on the CCR7 receptor during molecular dynamics simulation.

### 2.2 Recombinant product of exemplary cytokine based multi-epitope protein

Finally, to synthesize the gene construct, the final protein structure was translated and optimized based on the Ecoli bacterium, and the NheI and XhoI restriction site were placed at 5’ and 3’, respectively, and the KOZAK sequence was placed at 5’ to improve translation. The genescript company was constructed and inserted into the pET-28a vector for expression in the Ecoli bacterium. Then, specific primers of *nptII* gene was used to identify the presence of transgenes in bacteria by PCR. So, PCR was performed based on Parstous kit; Cat.No C101081. To study the expression of *ccl21* and *ccl19* genes in transgenic bacteria, Whole RNA was extracted from infiltrated leaf tissue (Parstous kit; Cat. No A101231) and complementary DNA (cDNA) was synthesized through reverse transcription using reverse primer (Parstous kit; Cat.No C101131). Also Real-time PCR (BioRad) was performed by using Parstous kit; Cat.No C101021. In order to purify the recombinant protein, a Ni-IDA Resin affinity column of Pars Toos Company was used. The NI-IDA nickel resin column is a protein purification device based on the presence of 6 histidine tags in the protein structure. The interaction between histidine and metal ions is the first step to protein purity. In this interaction, the histidine tag sequence is immobilized to the Ni2 + cation in the solid phase of the column using immunodiastic acid (IDA) groups. In this reaction, the impurities are removed and the purified proteins are dissolved by histidine tag by imidazole or by lowering the pH of the solution. Purified multiepitope protein was measured by standard protein dot blot assay(17). So 10μl of purified recombinant protein from transgenic strain was dotted on the nitrocellulose membrane, and it was incubated with BSA for 1 hour. After incubation, the membrane was thoroughly washed three times with PBST/PBS and incubated with conjugated anti-6x His tag® mouse monoclonal antibody for 1 hour at 37ºC, then washed three times with PBST/PBS, and finally incubated with DAB (diaminobenzidine) substrate. A small amount of commercial CCL21 antigen containing Histidine tag (1 μL) was used as the positive control (Cat No: 966-6C), and 10μl of protein from the wild type Ecoli was used as the negative control. For western blotting, the resulting SDS-PAGE was transferred to PVDF membrane by electroblotting (Biometra) for 2 hours at 120 V. The membrane was blocked with blocking buffer [BSA in PBS] for 1 hour at room temperature, Then it was cleansed three times with washing buffer [PBST]—each time 10 minutes at room temperature, then probed with the conjugated anti-6x His tag® mouse monoclonal antibody (Sigma-Aldrich) 1 hour at room temperature and washed three times with washing buffer (CMT; Cat.No CMGWBT). Finally, the protein bands were visualized by staining the membrane with TMB substrate (Promega; Cat.No W4121)(13). ELISA tests were performed to determine the amount of protein according to the existing protocol and ELISA reader. So, ELISA plates were coated with purified multiepitope protein and CCL21 antigen at 37°C for 1 hour, and was incubated with 1% BSA in PBS for 2 hours at a temperature of 37°C. The wells were washed using PBST/PBS, and incubated with the antiserum (conjugated anti-6x His tag® mouse monoclonal antibody), which was reactivated against commercia CCL21 (1:1000 dilutions) Wells were developed with TMB substrate. The color reaction was stopped by H2SO4 (2N concentration) and read at 405 nm of wavelength (CMT; Cat.No CMGTMB-100)(17).

### 2.3 Structural analysis of an exemplary cytokine based multiepitope protein

In the next step multi-epitope protein including purity were examined using MALDI-TOF/TOF mass spectroscopy, and also FTIR spectroscopy (35). In FTIR method, after drying the purified multiepitope protein extraction by using Freeze Dryer, dry samples were completely pressed to prepare a KBr tablet. So about 8.1 solids and 0.25-0.50 teaspoon full of KBr were thoroughly mixed in a mortar while grinding with the pestle. The material was pressed at 5000-10000 psi by pellet press instrument. The compressed sample was placed in the FTIR sample holder. Spectral analysis of UV absorption in the range 400-4000 cm-1 was performed by UNICAM UV 100 UV/visible light spectrophotometer(17). FTIR spectra were recorded on a Thermo Nicolet AVATAR370 spectrometer(35). MALDI TOF/TOF mass spectroscopy was used to illustrate no protein sequence of bacteria were found in this purified protein and just the epitope sequences of human genes were found in this multiepitope protein. So, the purified multiepitope protein was analyzed by SDS-PAGE and spot of protein were excised from stained gels, and Gel slices were distained with wash solution [100% acetonitrile and 50 mM ammonium bicarbonate (NH4CHO3)] for 1 hour at room temperature. The protein spot was then air-dried for 30 min at 37°C, then it was digested using a trypsin solution (12 ng/ml trypsin in 50 mM NH4CHO3) by incubation for 45 min at 47°C. Excess of trypsin solution was removed and 50 mM NH4CHO3 was replaced, then gel slice was incubated overnight at 37°C. Samples were sent to the Centre of Mass Spectrometry of York University(UK) for analysis of this recombinant multiepitope protein by conventional ionization methods. Finally, the results of MALDI TOF / TOF was received as a mgf file by YORK company and analyzed by Mascot software, the obtained sequences were blasted in NCBI database and their homology was examined(17).

### 2.4 Effect of cytokine based multi epitope protein on PBMC cells

First, the PBMC cells of healthy individuals and patients with colon and lung cancer were isolated from Musa Ibn Jafar Hospital in Mashhad,Iran and cultured. So whole blood samples of 30 healthy donors as control and 30 cancer patients (18 colon and 12 breast cancer samples) were collected and transferred to tubes containing anti-coagulant EDTA (it should be noted that T cells in PBMCs of individuals with CCR7+ receptor increase the binding affinity of CCL19 and CCL21 chemokines). Peripheral blood mononuclear cells (PBMC) were isolated by density gradient centrifugation according to manufacturer’s protocol. Isolated cells were cultured overnight in DMEM supplemented with 10% fetal bovine serum (FBS), 1% penicillin-streptomycin and 1% L-glutamine). More than 95% PBMC viability was obtained as evaluated by Trypan blue staining. Then, the effect of the exemplary cytokine-based multi-epitope protein on the PBMC cells was evaluated by incubating the PBMC cells with a commercial CCL21 or with the exemplary cytokine-based multi-epitope protein for 48 hours, and then RNA of the PBMC cells was extracted (Parstous kit; Cat. No A101231) for analyzing the gene expressions and cDNA was synthesized through reverse transcription using oligo(36) primer (Parstous kit; Cat.No C101131)(17). Real-time PCR (BioRad) was performed in a 20 μL reaction volume containing 1 μM of each primer and 10 μL of SYBR green real-time PCR master mix (Parstous kit; Cat.No C101021)(15). Real-time PCR experiments were carried out for each sample in duplicate. Forward and reverse primers for real-time PCR has been shown in Table1.

**Table 1:**
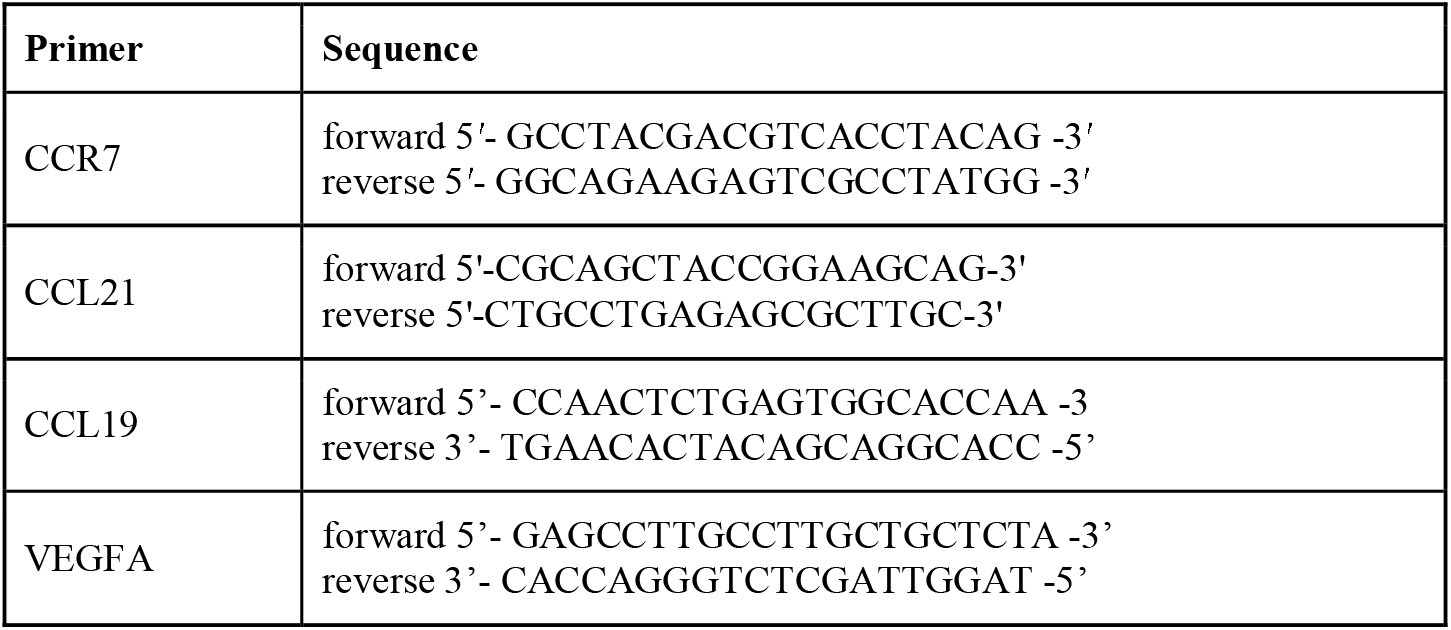
Forward and reverse primers for real-time PCR

### 2.5 Evaluation of cytotoxicity effect of cytokine based multi epitope protein

The MTT assay was used to determine the half-maximal inhibitory concentration (IC50) of the exemplary cytokine-based multi-epitope protein on the CCR7^+^ MCF7 cancer cell line. Three days after de-freezing and culturing the cells in DMEM medium using a CO2 incubator at 37°C, the cells were passaged and after reaching the appropriate density of cells, the toxicity of the cytokine-based multi-epitope protein on CCR7^+^ MCF7 cancer cells at 24, 48, and 72 hours after incubation using the MTT assay compared to the DMEM medium as a negative control group at different concentrations of 2.5, 5, 7.5, and 10 μg/ml, consistent with one or more exemplary embodiments of the present disclosure(15).

### 2.6 Evaluation of the effect of the multi epitope protein on cancer cell migration by Wound healing assay

The potential activity of the exemplary cytokine-based multi-epitope protein in stimulating the proliferation and migration of cancer cells was investigated using the wound healing assay. Clinical studies have discovered that MCF7 breast cancer tumors express CCR7. MCF7/CCR+ cells were obtained from Academic Center for Education Culture and Research (ACECR) of Khorasan Razavi Branch, Mashhad, Iran and were seeded in 6-well plates at densities of 6×150 cells/well in the DMEM high glucose growth medium. After 95% confluency, cells that grew as a monolayer was scratched using a sterile pipetting tip, drawn firmly across the dish to induce in-vitro wounds, Cells were washed by PBS to remove the loosened debris. The initial width of scratches in cancer cell cultures was estimated between 700 to 800 μm. In the next step, 7.5 μg/ml of the exemplary cytokine-based multi-epitope protein, commercial CCL21 protein as the positive control, and DMEM medium as the negative control were added to a set of each well, Moreover, the cells were treated with an equivalent amount of DMSO, as control. For migration assay, cultures were rinsed twice using PBS solution, fixed by absolute methanol and stained by Giemsa. Images were recorded at 24, 48, and 72 hours after wounding. The rate of cell migration and closure of the scratch were quantified by evaluating the changes in the wound area (pixels) using Image J software(14).

### 2.7 Evaluation of recombinant protein on the migration of immune cells by Chemotaxis assay

Chemotaxis assay was used to investigate the effect mechanism of CCL21 and CCL19 of multiepitope protein in the migration of T-cells, since chemokines that are involved in leukocyte migration(17). So, to create wells on the agarose gel, 1% agarose was prepared in a medium composed of 50% DMEM, 10% FBS, 50% PBS, and 2 mM L-glutamine. 1.6 mL of agarose solution 1% was added to each 10-mm sterile petri dish. To humidify the gel, 5 mL DMEM was added to each petri dish after 20 min of cooling the gel. Afterward, 5 mL FBS free DMEM was added to the gel for 1–6 hours before performing the cell migration assay, after which three small wells with a distance of 10 mm were designed in the Petri dishes. PBMC cells were seeded at a density of 1 × 10^6^ cells in the middle well with 10% FBS-DMEM. After 24 hours, the medium was replaced by FBS-free DMEM. Due to their chemoattractant activity, chemokines (commercial CCL21 and recombinant multiepitope protein) were then placed in one of the neighboring wells, with FBS-free DMEM or non-recombinant protein in other wells as the negative control. Image capture and measurements were accomplished using an invert microscope (Olympus). For DAPI staining, cell pellets were fixed in 4% paraformaldehyde for 8 minutes (Sigma, Germany). For washing the cells, each well was washed with 60 μL of 1x PBS three times for 5 minutes. Afterward, the cells were permeabilized with 60μL/well of 0.1% Triton X-100 for 10 minutes and stained with 50μL/well of DAPI (1:2000 dilution, in 1x PBST) (Merck, Germany) for 10 min. The stained cells were then counted using a fluorescent microscope. Chromatin condensation and nuclear fragmentation were the criteria to confirm apoptosis(17).

### 2.8 Animals and experimental design

Twenty female BALB/c mice, 6–8 weeks of age were obtained from Ferdowsi University of Mashhad (Mashhad, Iran). During the entire experimental period, animals were kept in colony cages in a pathogen-free and controlled conditions with usual light/dark schedule (light/ dark cycle of 12/12 hours). All mice were acclimated for one week before the investigation and adequately served with standard chow diet and water ad libitum. The experimental protocols of present study were approved by the local ethics committee on animal care (Mashhad University of Medical Sciences, approval ID: IR.MUMS.RES.1399.428). Following adaptive feeding, mice were assigned to four groups. Five healthy mice in control group (n=5) were intraperitoneal injected with 150 μg physiological normal saline (Group A) and five mice injected with 150 μg recombinant antigen (Group B) whereas five mice in the tumor group (n=5) received 150 μg recombinant antigen (Group C) also five tumor mice injected with 150 μg physiological normal saline (Group D). After 48h, animals were anesthetized intraperitoneal and their blood samples collected through cardiac puncture. For this purpose, the animals were fixed, and blood was collected from the heart using 8 cc syringe. Then the blood samples were centrifuged at 3000 rpm for 18 minutes to separate the serum. The collected serum samples were kept at -20°C until biochemical parameters were measured. In order to measure white cell counts, the blood samples taken in all groups only be done in CBC vials containing EDTA anticoagulant (Heparin tubes not recommended as an anticoagulant for cell counts, because the cells clump in heparin, invalidating counts), and all tubes were placed in the hematology roller mixer for 29 minutes to mix the samples. Then the samples were given to the Sysmax model tuberculosis counter hematology machine and the white blood cells as well as Neutrophils, Basophils, Monocytes, Lymphocytes and Eosinophils were measured. The collected data was tested by Kolmogorov-Smirnov test and this is appearing that these data is not Normal. So, for comparing the mean rank of groups we used the Mann-Whitney Test. The result shown that there is significant difference between some groups.

## 3 Result and Discussion

### 3.1 Design and Insilico analysis of cytokine-based multi-epitope protein

Cytokine-based multi-epitope protein including different epitopes of human Complex II and I (MHCII/MHCI) (37) and CCR7 receptor, which is present on different immune cells, including T helper lymphocytes (THL), cytotoxic T lymphocytes (CTL) and B lymphocytes. The CCL21 and CCL19 sequences selected epitopes had a DCCL motif, a domain binding to the CCR7 receptor, and a Pan HLA DR-binding epitope (PADRE) peptide sequence and a putative glycosaminoglycan binding site that covers more than 90% of the HLA alleles. CCL21 and CCL19 are chemokines that control cell trafficking and are involved in numerous pathologic and inflammatory conditions, and it is endocytosed with its receptor (CCR7), via both MHC class I and II processing pathways to induce CD8^*+*^ and CD4^+^ T-cell responses. These data suggest that the exemplary cytokine-based multi-epitope protein can be taken up, processed, and presented by APCs after binding and internalization through the CCR7 receptor. CCR7 facilitates the uptake and processing of tumor antigens to induce efficient CD4^+^ T-cell responses both in-vitro and in-vivo using the MHC class II antigen processing pathway. Since the selected epitopes of CCL21 and CCL19 have a half-maximal inhibitory concentration (IC50) less than 50, they can bind to both MHCI and MHCII molecules. In addition, a part of selected epitopes of IL-1β is involved in inflammatory and immune responses and has high adjuvant effects. While the selected epitope of GMCSF adjuvant covers more than 90% of HLA alleles and can only bind to MHCI molecules, it does not affect MHCII molecules since its IC50 was greater than 50. By binding the cytokine-based multi-epitope protein to MHCI and MHCII molecules in T-cells and CCR7 receptors, the cytokine-based multi-epitope protein can produce anti-metastatic and cytotoxicity effects on cancer cell lines and chemotactic response in lymphocyte cells. The cytokine-based multi-epitope protein may be recognized as endogenous chemokine via its CTL cell epitopes that can bind to MHCI. Also, binding cytokine-based multi-epitope protein to MHC II may occur via the exogenous pathway of antigen presentation, which activates T helpers. The gene construct also contained rat chemokine KC as a signal peptide and a polyhistidine tag for purification of the gene construct. The exemplary cytokine-based multi-epitope protein has an instability index of about 30.5, which indicates the stability of the recombinant multi-epitope protein. Also, the aliphatic index of the exemplary cytokine-based multi-epitope protein is about 84.57, which indicates the temperature resistance of the exemplary cytokine-based multi-epitope protein. Moreover, the grand average of hydropathicity of the exemplary cytokine-based multi-epitope protein is about -1.25, which indicates that the vaccine is hydrophilic. The exemplary cytokine-based multi-epitope protein has a solubility index of about 84.3%. The in-vitro half-life of the exemplary cytokine-based multi-epitope protein is less than about 30 hours in mammalian cells, less than about 20 hours in yeasts, and less than about 10 hours in *Escherichia coli*. Furthermore, the allergenicity of the cytokine-based multi-epitope protein was evaluated, and it was shown that this protein has 98% non-allergenicity. Therefore, this recombinant multi-epitope protein can be used for treatment purposes.

### 3.2 Molecular dynamic simulation of cytokine-based multi-epitope protein

In this modeling method, 3D structures with sequences very similar to the target sequence are used as the model. Ten models were produced during the modeling step, and one with the most stability, interaction, and connectivity between the epitopes of the exemplary cytokine-based multi-epitope protein was selected as the best model (FIG. 1). FIG. 1 illustrates a three-dimensional (3D) structure 100 of an exemplary cytokine-based multi-epitope protein, consistent with one or more exemplary embodiments of the present disclosure. Referring to FIG. 1, the exemplary cytokine-based multi-epitope protein may include different epitopes of different proteins, including IL-1β and signal peptide 102, GM-CSF 104, CCL19 106, and CCL21 108. illustrates RMSD changes related to exemplary cytokine-based multi-epitope protein during 100 nm of molecular dynamics simulation, consistent with one or more exemplary embodiments of the present disclosure. Referring to FIG. 2, at the beginning of the MD simulation, the RMSD diagram shows an uptrend. In the first 10 nanoseconds of the simulation, the slope of the increase in RMSD is so rapid that after about 10,000 PCM, the RMSD value reaches 1 nm, but the increase slows down; so that at 60,000 pm, the RMSD value is equal to 25/1 nanometer. After this time, the RMSD value decreased slightly and reached 1.1 nm at 70,000 picoseconds, and remained constant at the end of the simulation, indicating the stability of the protein structure at the end of the simulation. The radius of Gyration is one of the critical parameters in studying changes in protein size during the MD simulation. The lower the radius of Gyration during MD simulation, the more compact the protein. Conversely, as the radius of Gyration increases, the size of the protein increases too. FIG. 3 illustrates some changes in the radius of Gyration of an exemplary cytokine-based multi-epitope protein during 100 nm of molecular dynamics simulation, consistent with one or more exemplary embodiments of the present disclosure. Referring to FIG. 3, the value of the radius of Gyration of an exemplary cytokine-based protein equals about 2.62 nm at the beginning of the MD simulation. However, as the simulation continues, the value of the radius drops rapidly and reaches 2.29 nm in about 10,000 picoseconds. From this time onward, the change in the slope of the radius of Gyration slows down, such that at 80,000 picoseconds, the value is equal to about 2.21 nm, after which it remains constant until the end of the simulation which is indicating stability. FIG. 4A illustrates the 3D structure of an exemplary multi-epitope protein before 100 nanoseconds of MD simulation, consistent with one or more exemplary embodiments of the present disclosure. FIG. 4B illustrates the 3D structure of an exemplary cytokine-based multi-epitope protein after 100 nanoseconds of MD simulation, consistent with one or more exemplary embodiments of the present disclosure. Referring to FIGs. 4A-4B, the 3D structure of cytokine-based multi-epitope protein is shown from two sides and top angles. Equilibration was then performed under NPT conditions with a time step of 2 femtoseconds for 4 nanoseconds. The Berendsen algorithm was used to keep the temperature constant at 310 K and the pressure at 1 atmosphere in NVT and NPT conditions. Also, during MD simulation, the RMSD parameter was used to study the stability of the exemplary cytokine-based multi-epitope protein on the CCR7 receptor during the MD simulation. FIG.5 illustrates a molecular docking complex between cytokine-based multi-epitope protein and CCR7, consistent with one or more exemplary embodiments of the present disclosure.

**FIG. 1.**
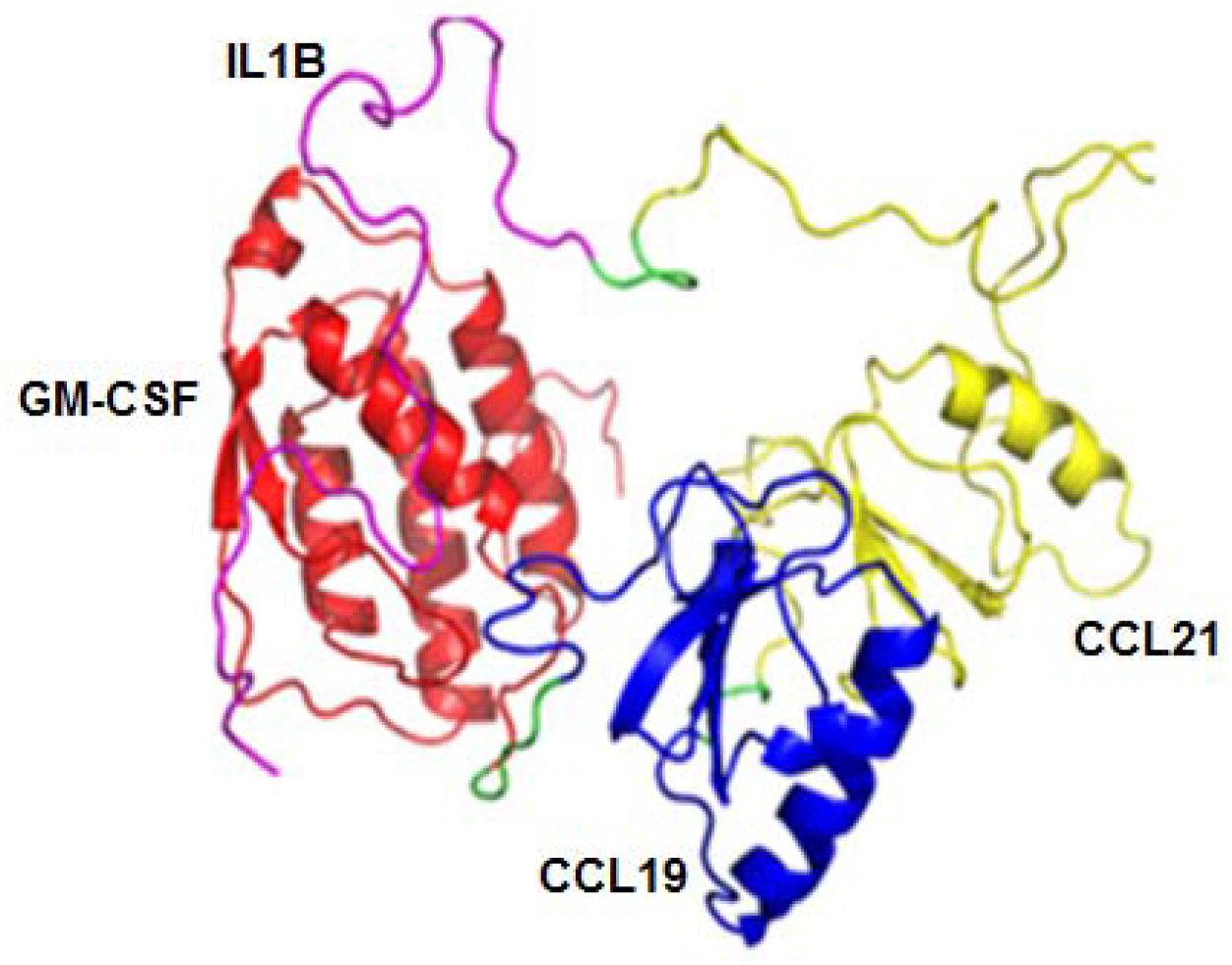
illustrates a three-dimensional (3D) structure of an exemplary cytokine-based multi-epitope protein, consistent with one or more exemplary embodiments of the present disclosure.

**FIG. 2.**
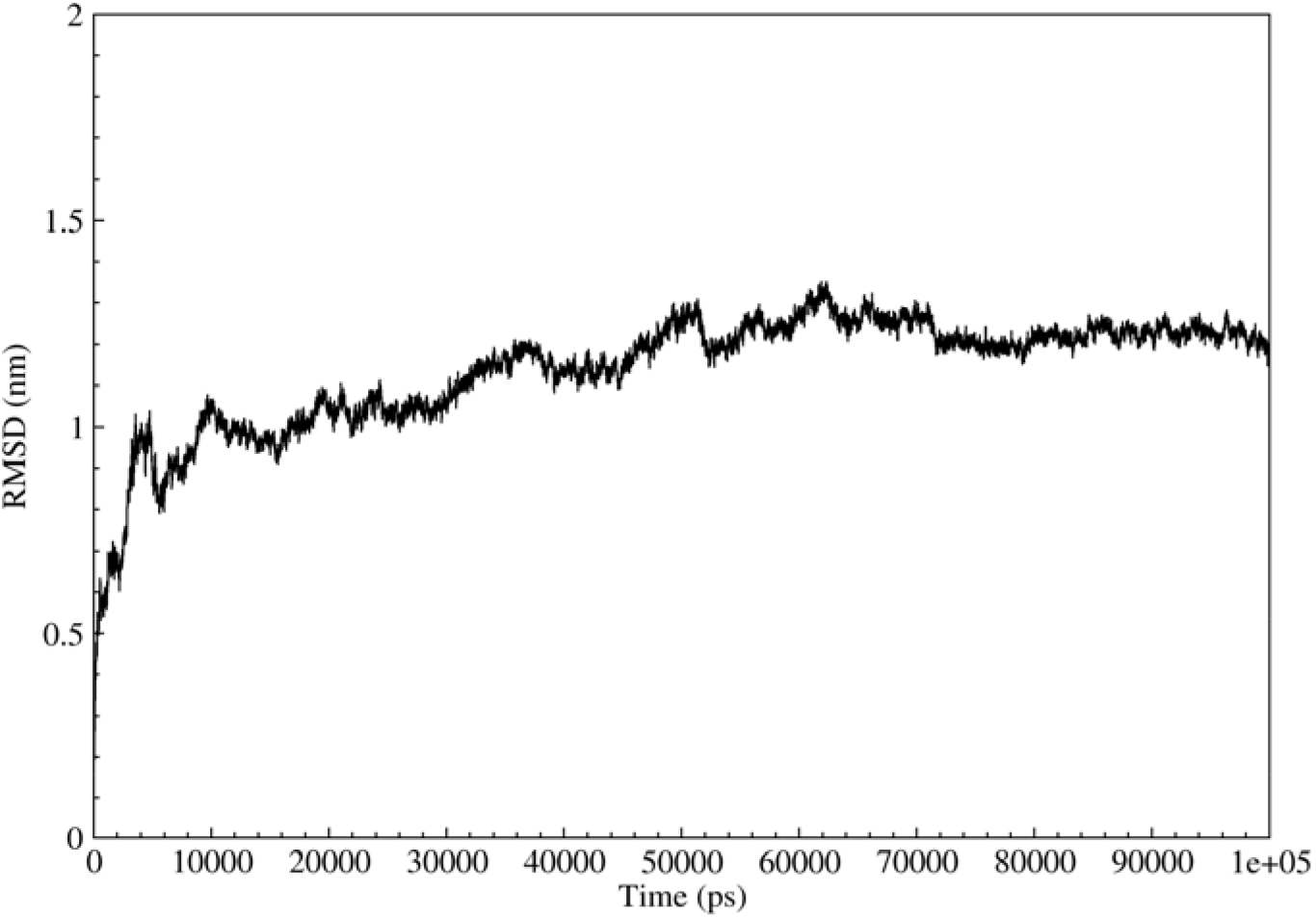
illustrates root deviation of the mean squares (RMSD) changes related to exemplary cytokine-based multi-epitope protein during 100 nm of molecular dynamics simulation, consistent with one or more exemplary embodiments of the present disclosure.

**FIG. 3.**
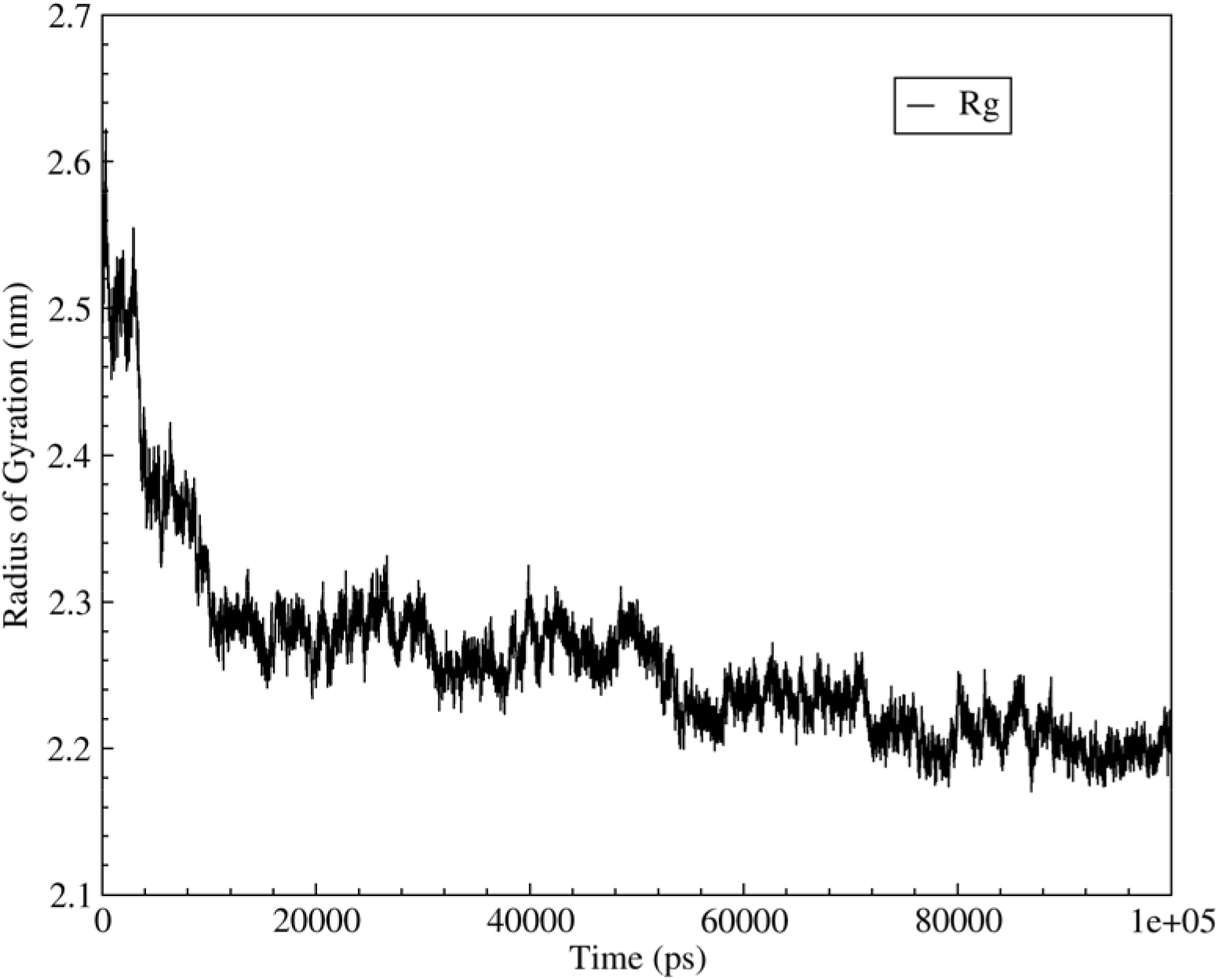
illustrates changes in the radius of Gyration of an exemplary cytokine-based multi-epitope protein during 100 nm of molecular dynamics simulation, consistent with one or more exemplary embodiments of the present disclosure.

**FIG 4.**
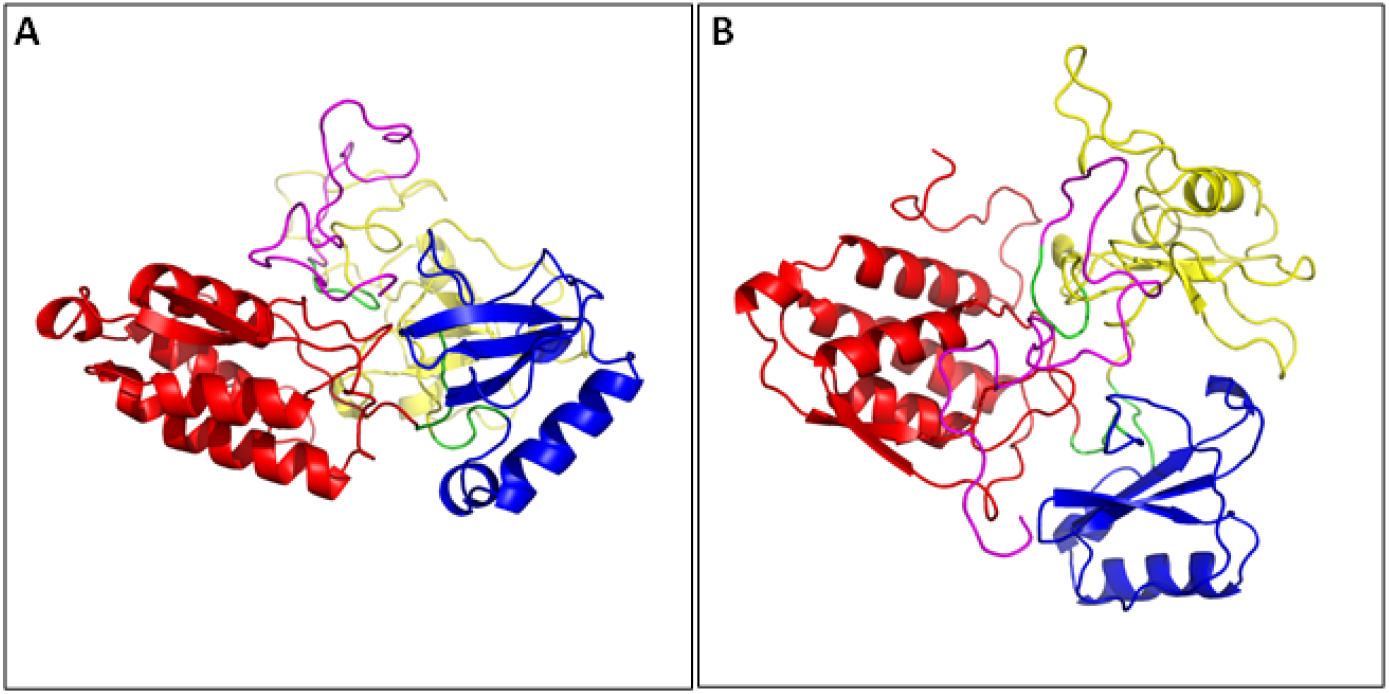
The 3D structure of the exemplary cytokine-based multi-epitope protein. **FIG. 4A** illustrates the 3D structure of an exemplary multi-epitope protein before 100 nanoseconds of MD simulation, consistent with one or more exemplary embodiments of the present disclosure. **FIG. 4B** illustrates the 3D structure of an exemplary cytokine-based multi-epitope protein after 100 nanoseconds of MD simulation, consistent with one or more exemplary embodiments of the present disclosure.

FIG. 6 Shows the change of RMSD diagram of the exemplary cytokine-based multi-epitope protein in the CCR7 binding state during MD simulations, consistent with one or more exemplary embodiments of the present disclosure. Referring to FIG. 6, the value of RMSD shows a sharp increase at the beginning of the simulation, and after a time of 70,000 picoseconds, it reaches 0.8 nm, and after this time until the end of the simulation, it remains on the same relative stability value. Also, the binding power of exemplary cytokine-based multi-epitope protein to the CCR7 receptor was evaluated using the molecular mechanics Poisson-Boltzmann surface area (MMPBSA) method, So van der Waals and electrostatic energies have the same effect on the connection power and play a more significant role in the connection power. Also, the amount of polar solvent energy is equal to 142.388. Finally, the amount of free energy binding of the exemplary cytokine-based multi-epitope protein to the CCR7 receptor obtained from this calculation equals -60.592 kg/mol. The more negative the bond energy, the greater the bond strength.

**FIG. 5.**
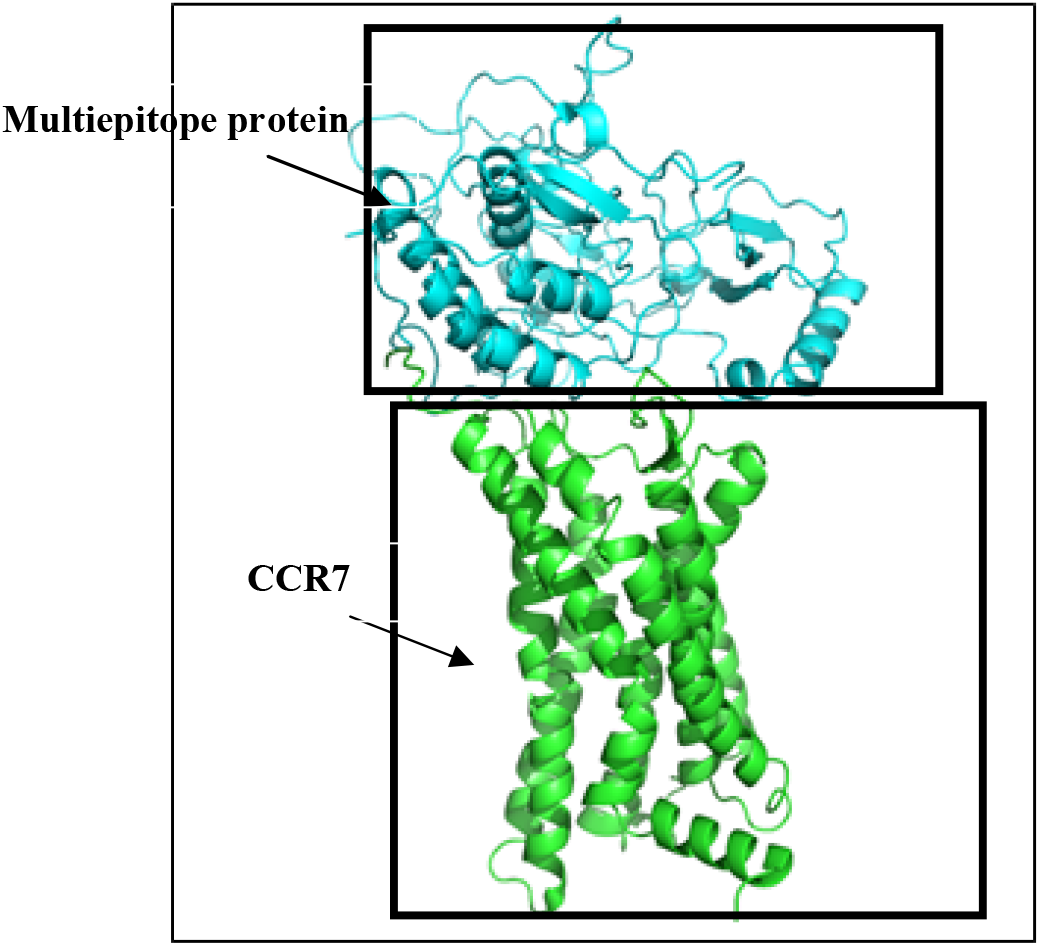
illustrates a molecular docking complex between an exemplary cytokine-based multi-epitope protein and CCR7, consistent with one or more exemplary embodiments of the present disclosure.

**FIG. 6.**
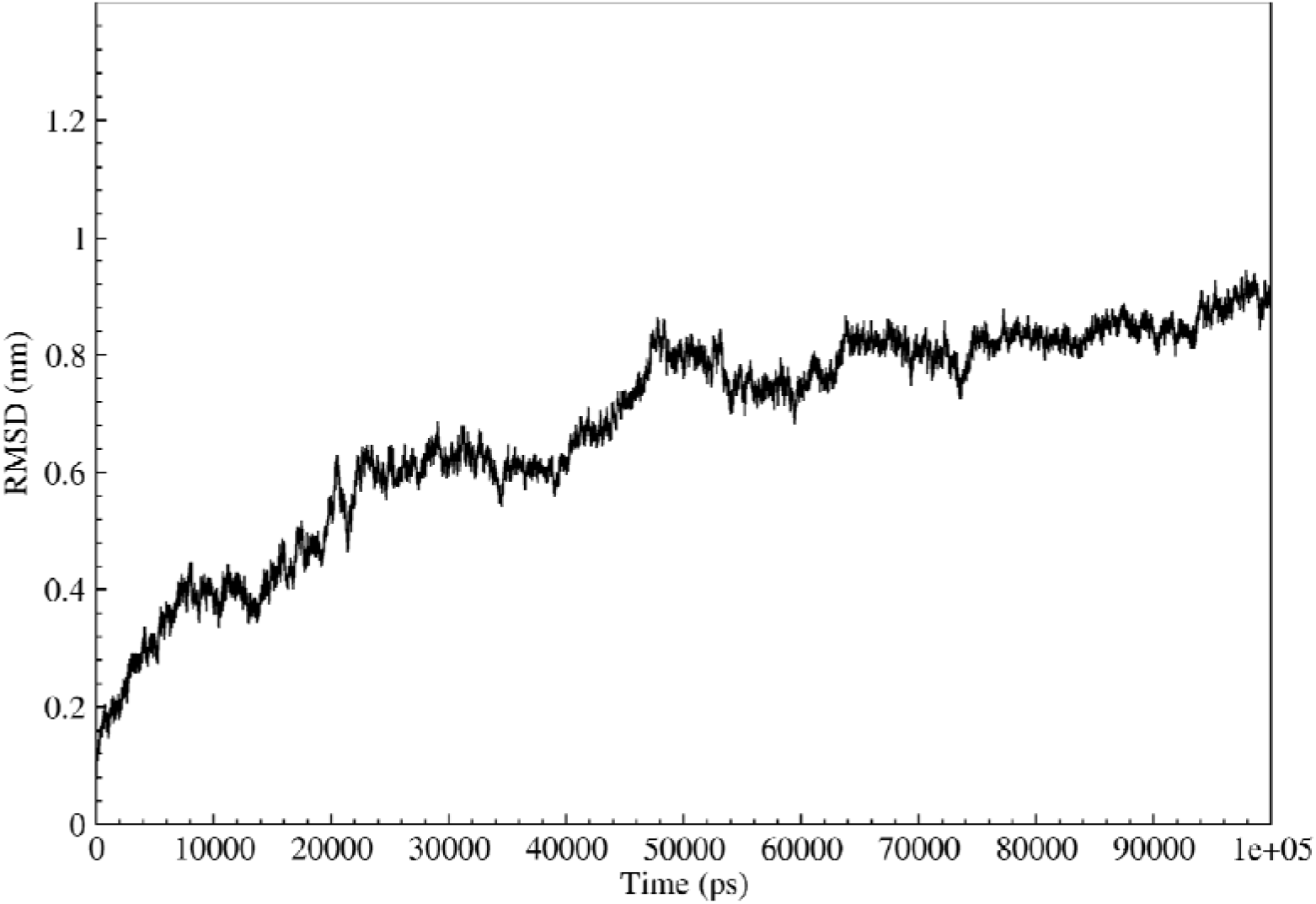
illustrates changes in the RMSD diagram of the exemplary cytokine-based multi-epitope protein in the CCR7 binding state during MD simulations, consistent with one or more exemplary embodiments of the present disclosure.

### 3.3 Recombinant production of cytokine based multi-epitope protein

The recombinant expression of cytokine-based multi-epitope protein in *E. coli* was studied by real-time PCR. FIG. 7 illustrates relative expression of CCL21 and CCL19 epitopes in *E. coli* compared to the beta-actin gene as a housekeeping gene using real-time PCR, consistent with one or more exemplary embodiments of the present disclosure. Referring to FIG. 7, the expression of CCL21 and CCL19 epitopes is about 2.5-fold and 3-fold higher than that of the beta-actin gene. NC group relates to results of non-transgenic *E. coli* as a negative control. The purified cytokine-based multi-epitope protein expression was assessed by sodium dodecyl sulfate-polyacrylamide gel electrophoresis (SDS-PAGE), dot-blot, and Western-blot assays. FIG. 8A, a 65 kDa band is observed in well 6 for fractions resulting from elution with 250 mM imidazole buffer solution, which indicates the molecular weight of the exemplary cytokine-based multi-epitope protein. On the other hand, there is no 65 kDa band in well 7 of non-transgenic E. coli, indicating no transgene expression in non-transgenic bacteria. In an exemplary embodiment, the cytokine-based multi-epitope protein may have a molecular weight between about 60 kDa and about 65 kDa. In an exemplary embodiment, exemplary cytokine-based multi-epitope protein may further include a purification tag, including at least one of a polyhistidine tag and a glutathione S-transferase (GST) tag. FIG. 8B illustrates Western blot results of an exemplary cytokine-based multi-epitope protein including well M: a molecular marker of protein, well 1 and well 2: exemplary cytokine-based multi-epitope protein, well 3: commercial CCl21 antigen, and well 4: Negative control (total protein extracted from non-transgenic *E. coli*), consistent with one or more exemplary embodiments of the present disclosure. Referring to FIG. 8B, the 65 kDa band is visible in the recombinant protein extracted from the transgenic bacteria in wells 1 and 2. The band of about 40 kDa corresponds to the complete sequence of CCL21 commercial antigen as positive control is visible in well 3. Also, there is no observation of protein band in well 4 of total non-transgenic bacterial protein as the negative control. Moreover, an enzyme-linked immunosorbent assay (ELISA) was performed to determine the amount of exemplary cytokine-based multi-epitope protein using an antibody against the polyhistidine tag. FIG. 9 illustrates quantitative measurement of exemplary cytokine-based multi-epitope protein using ELISA, including a transgenic bacteria (TG) expressing the exemplary cytokine-based multi-epitope protein, non-transgenic bacteria as a negative control (NC), and bovine serum albumin (BSA) as a blank group, consistent with one or more exemplary embodiments of the present disclosure. Referring to FIG. 9, the transgenic bacteria group (TG) has the maximum and consistent expression of the exemplary cytokine-based multi-epitope protein, about 2.14% of total serum protein (TSP). Due to non-specific reactions, the observed signal in wild-type and BSA samples can be ignored.

**FIG. 7.**
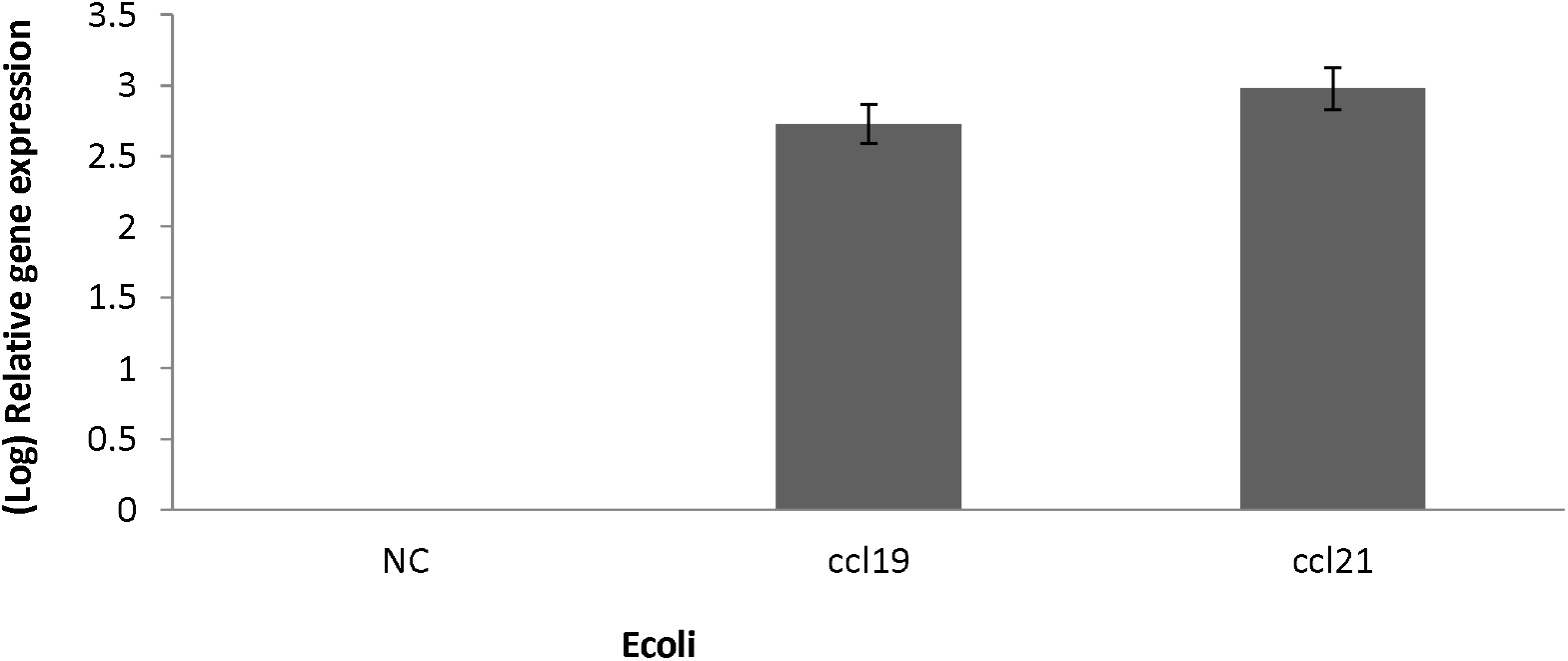
illustrates relative expression of CCL21 and CCL19 epitopes in *E. coli* compared to the beta-actin gene as a housekeeping gene using real-time PCR, consistent with one or more exemplary embodiments of the present disclosure.

**FIG. 8A.**
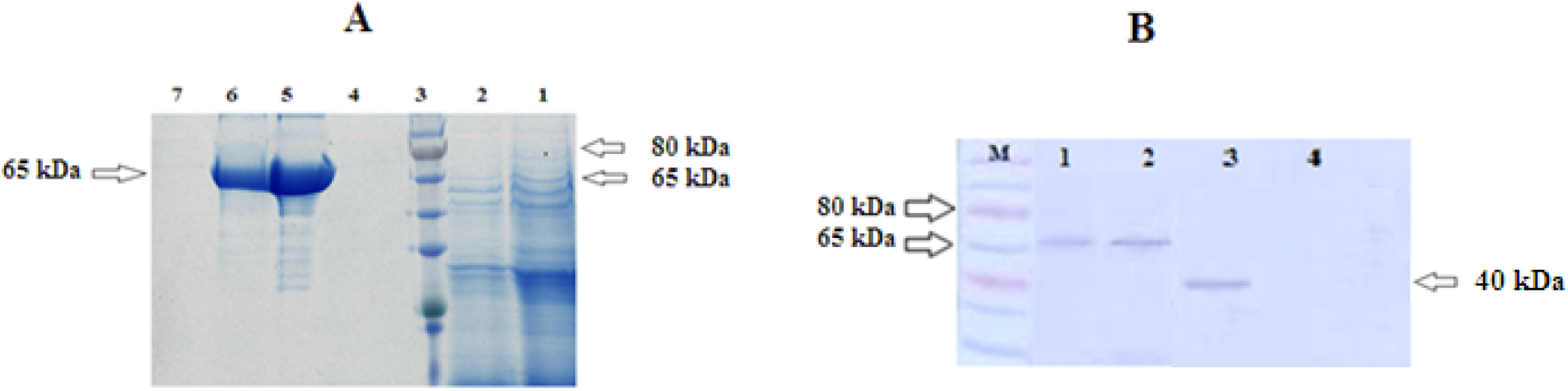
illustrates an image of SDS-PAGE of exemplary cytokine-based multi-epitope protein, including well 1: total protein extracted from transgenic *E. coli*, well 2: fraction flowing from the column, well 3: protein ladder, well 4: fractions resulting from washing with 500 mM imidazole buffer solution, well 5: fractions resulting from rinsing with 100 mM imidazole buffer solution, well 6: fractions resulting from elution with 250 mM imidazole buffer solution, well 7: Negative control (total protein extracted from non-transgenic *E. coli*), consistent with one or more exemplary embodiments of the present disclosure. **FIG. 8B** illustrates Western blot results of an exemplary cytokine-based multi-epitope protein including well M: a molecular marker of protein, well 1 and well 2: exemplary cytokine-based multi-epitope protein, well 3: commercial CCl21 antigen, and well 4: Negative control (total protein extracted from non-transgenic *E. coli*), consistent with one or more exemplary embodiments of the present disclosure.

**FIG 9.**
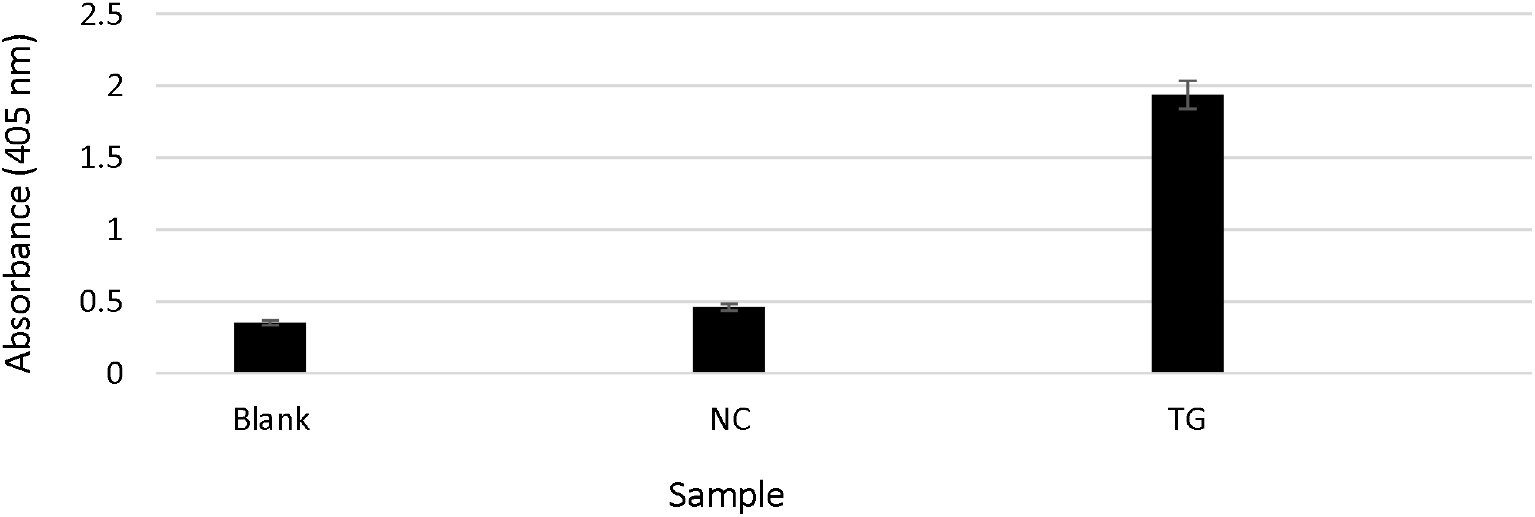
illustrates quantitative measurement of exemplary cytokine-based multi-epitope protein using ELISA, including a transgenic bacteria (TG) expressing the exemplary cytokine-based multi-epitope protein, non-transgenic bacteria as a negative control (NC), and bovine serum albumin (BSA) (blank), consistent with one or more exemplary embodiments of the present disclosure.

### 3.3. Structural analysis of cytokine based multiepitope protein

MALDI-TOF/TOF results were analyzed using the Mascot server. According to the MALDI-TOF/TOF mass spectroscopy, it was shown that EHVNAIQEARRLLNLSR and RLQRTSAKMKR as a parts of human CCL19 have molecular weight of 2020Da and 1380 Da respectively; SIPAILFLPRKRSQAE, VQEESNDK and EMFDLQEPTCLQT sequences are part of human CCL21, ILIβ and GM CSF with molecular weights of 1830 Da, 947.1 Da and 1560 Da respectively, whereas no bacteria sequences were detected. The average protein sequence coverage confirmed recombinant protein as the target protein that was correctly expressed and purified from transgenic bacteria. As a result, no protein sequence of bacteria was found in this purified protein, and only the epitope sequences of the exemplary cytokine-based multi-epitope protein were found.

Also, the structure and post-translation modification of the exemplary cytokine-based multi-epitope protein were analyzed by infrared spectroscopy using Fourier-transform infrared spectroscopy (FTIR) spectrophotometer. FIG. 10 illustrates Fourier-transform infrared spectroscopy (FTIR) spectra of the exemplary cytokine-based multi-epitope protein and commercial CCL21 antigen (B), consistent with one or more exemplary embodiments of the present disclosure. Referring to FIG. 10, the amine, glycoside, and phospholipid factor groups were examined by studying the peaks, and similar peak positions were observed in the spectra of the exemplary cytokine-based multi-epitope protein) and commercial CCL21 antigen as control. There is a peak difference in the 3000 cm^-1^-3500 cm^-1^ region, which is related to the presence of OH and NH2 groups in the commercial CCL21 antigen (B). Also, there is another peak difference in the absorption region of the CO carbonyl group (1500-2000 cm^-1^). Also the present of transmittance of commercial CCL21 was 68% but cytokine-based multi-epitope protein was 80%. Assessments of the FTIR analysis and MALDI-TOF/TOF mass spectrometry displayed that the exemplary cytokine-based multi-epitope protein was correctly expressed in *E. coli*.

**FIG. 10.**
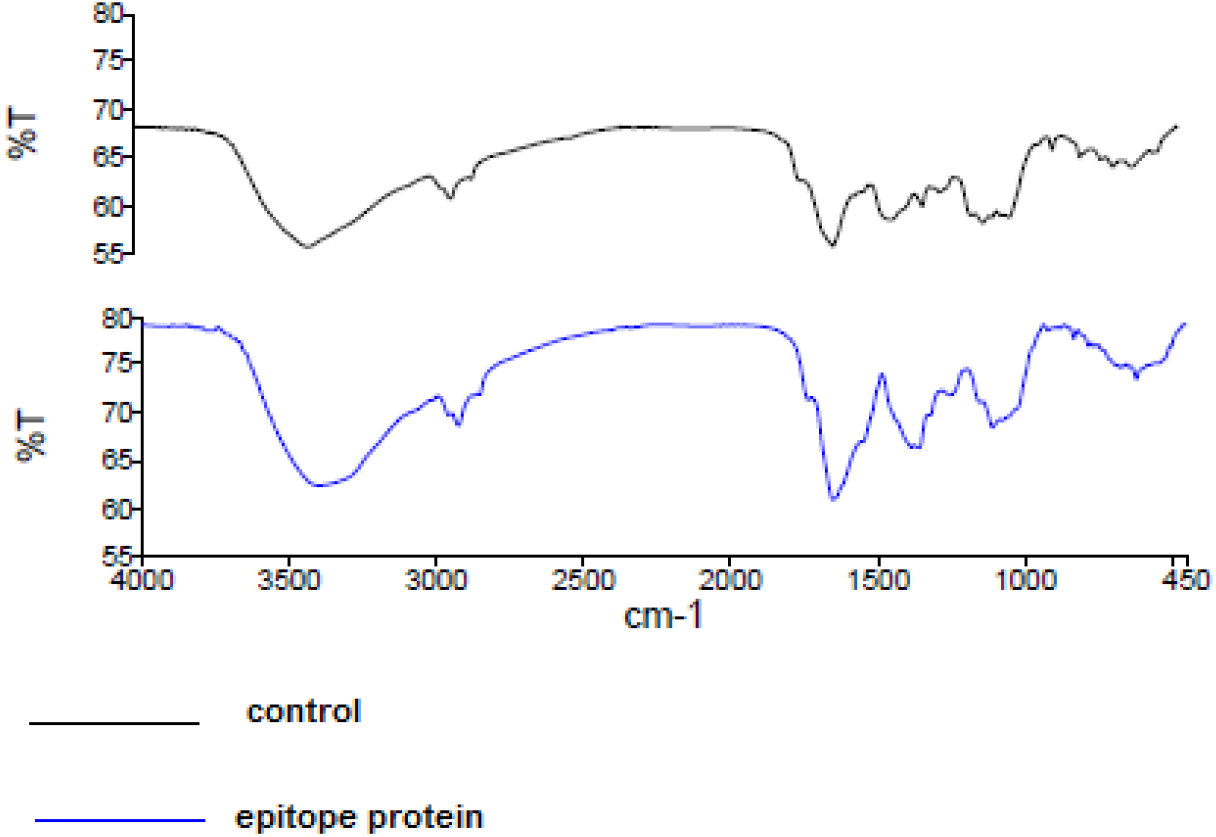
illustrates Fourier-transform infrared spectroscopy (FTIR) spectra of the exemplary cytokine-based multi-epitope protein (A) and commercial CCL21 (B), consistent with one or more exemplary embodiments of the present disclosure

### 3.4 Functional effect of cytokine based multiepitope protein

While an exemplary cytokine-based multi-epitope protein has an immunostimulatory effect, it may increase the expression of different immune genes, including CCL19, CCL21and CCR7 in PBMC cells after incubating with an exemplary cytokine-based multi-epitope protein. In this example, expressions of CCL19, CCL21, and CCR7 genes in the PBMC cells after incubation with an exemplary cytokine-based multi-epitope protein and commercial CCL21 as a positive control were studied by real-time PCR. FIG. 11 illustrates relative gene expressions in different groups, including PBMC cells with only DMEM medium as a negative control (NC), expression of CCR7 (A), CCL19 (B), and CCL21 (C) in PBMC cells of cancer samples, expression of CCR7 (D), CCL19 (E), and CCL21 (F) in PBMC cells of healthy samples, expression of CCR7 (G), CCL19 (H), and CCL21 (I) in PBMC cells of cancer samples treated with cytokine-based multi-epitope protein, expression of CCR7 (J), CCL19 (K), and CCL21 (L) in PBMC cells of cancer samples treated with commercial CCL21, consistent with one or more exemplary embodiments of the present disclosure.

**FIG. 11.**
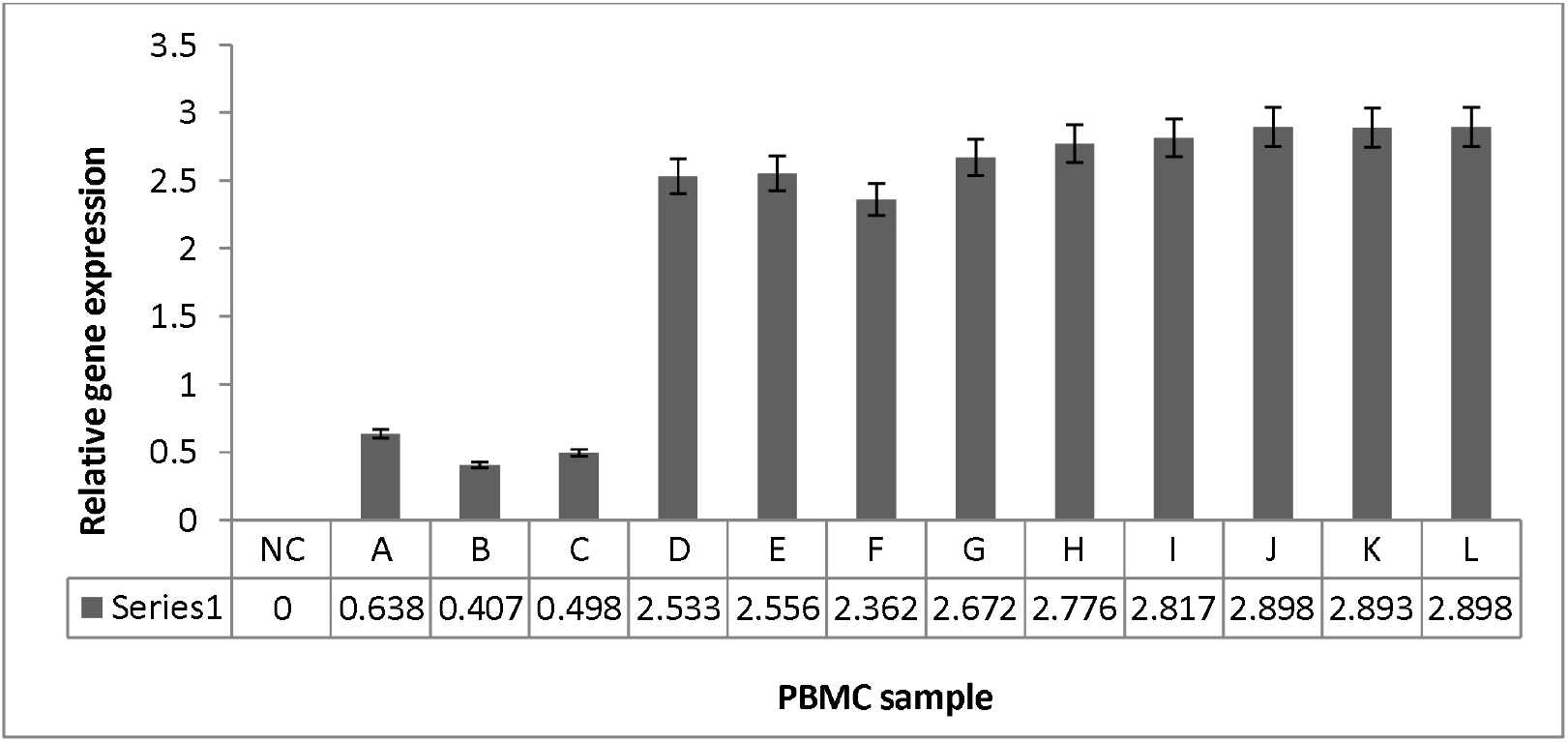
illustrates relative gene expressions in different groups, including DMEM medium without PBMC cells as a negative control (NC), expression of CCR7 (A), CCL19 (B), and CCL21 (C) in PBMC cells of cancer samples, expression of CCR7 (D), CCL19 (E), and CCL21 (F) in PBMC cells of healthy samples, expression of CCR7 (G), CCL19 (H), and CCL21 (I) in PBMC cells of cancer samples treated with cytokine-based multi-epitope protein, expression of CCR7 (J), CCL19 (K), and CCL21 (L) in PBMC cells of cancer samples treated with commercial CCL21, consistent with one or more exemplary embodiments of the present disclosure.

Referring to FIG. 11, the results showed that the expressions of CCR7, CCL19, and CCL21 genes in the group treated with the exemplary cytokine-based multi-epitope protein and treated with the commercial CCL21 antigen as the positive control were higher than that of the group with only DMEM as the negative control. Also, the expression of CCR7, CCL19, and CCL21 genes in cancer patients is higher than in a treated sample; therefore, the expression of these genes can be used as a biomarker for the early detection of cancers. It should be noted that T-cells in PBMCs of individuals with CCR7^+^ receptors increase the binding affinity of CCL19 and CCL21 chemokines to the CCR7 receptors.

### 3.4 Cytotoxicity assay of cytokine based multiepitope protein

Cytotoxicity of the exemplary cytokine-based multi-epitope protein was assessed. FIG. 12A illustrates an evaluation of the toxicity of the exemplary cytokine-based multi-epitope protein on CCR7^+^ MCF7 cancer cells at 24, 48, and 72 hours after incubation using the MTT assay compared to the DMEM medium as a negative control group at different concentrations of 2.5, 5, 7.5, and 10 μg/ml, consistent with one or more exemplary embodiments of the present disclosure. Referring to FIG. 12A, the survival rate of cancer cells incubated with purified recombinant protein at a 7.5 μg/ml concentration after 72 hours was 29.5% compared to the DMEM culture medium as control group. Also, the IC50 was 2.8 μg/ml. FIG. 12B illustrates an evaluation of the toxicity of a commercial CCL21 antigen on MCF7 cancer cells at 24, 48, and 72 hours after incubation using the MTT assay compared to the negative control group at different concentrations of 2.5, 5, 7.5, and 10 μg/ml, consistent with one or more exemplary embodiments of the present disclosure. Referring to FIG. 12B, the results showed that at a 7.5 μg/ml concentration, the survival of cancer cells incubated with commercial CCL21 protein was 33% compared to the DMEM medium as the control group. In examining the hypothesis of “significant difference between concentrations (7.5 with other concentrations) in the 72-hour period”. So we performed the ANOVA statistical test with Sheffe’s within-group test for recombinant, non-recombinant and commercial proteins separately. Since the P-Value for comparing (2.5 and 7.5), (5 and 7.5) and (10 and 7.5) was less than 0.05, therefore the hypothesis is accepted.

**FIG. 12.**
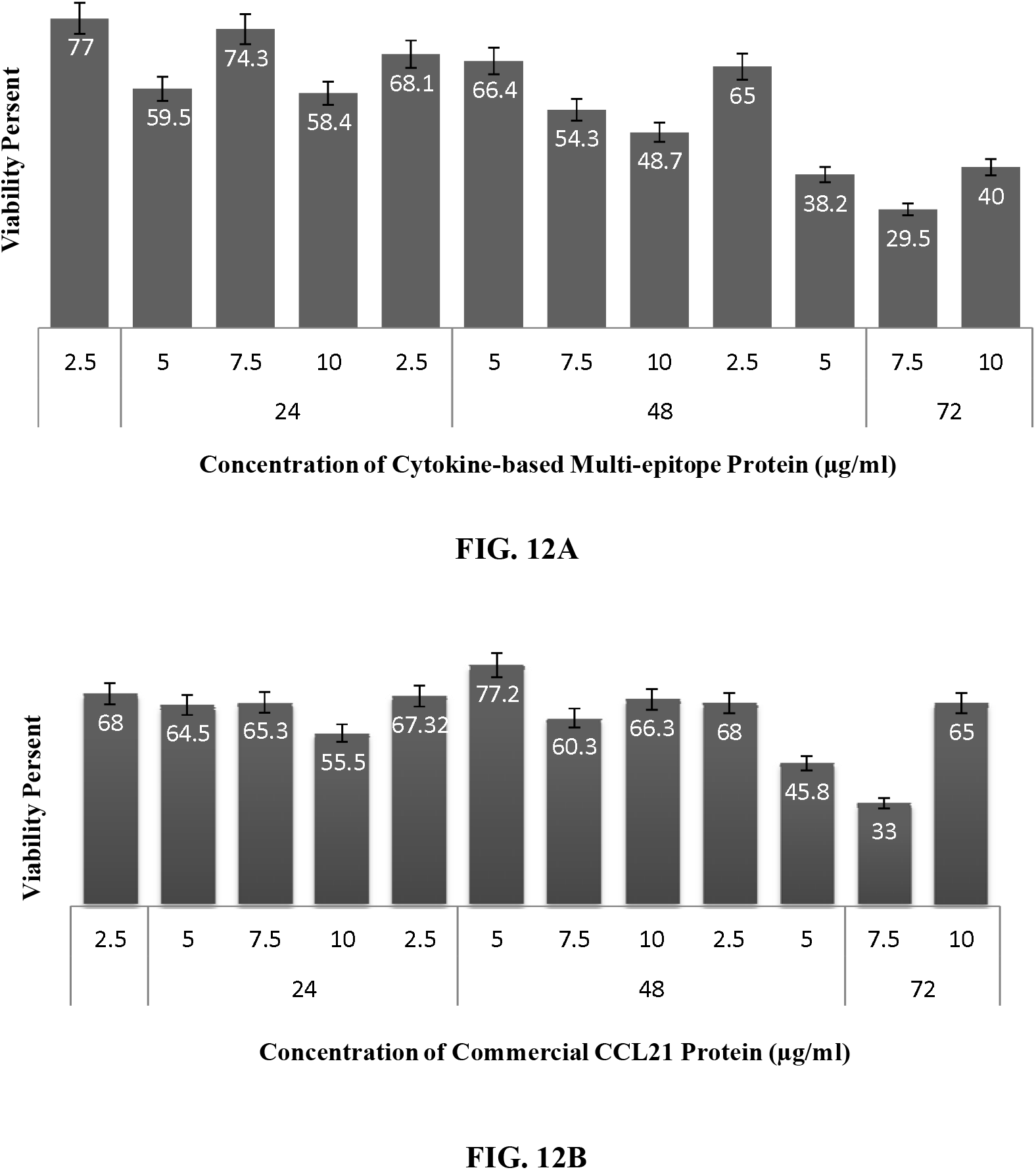
A illustrates an evaluation of the toxicity of the exemplary cytokine-based multi-epitope protein on CCR7^+^ MCF7 cancer cells at 24, 48, and 72 hours after incubation using the MTT assay compared to the DMEM medium as a negative control group at different concentrations of 2.5, 5, 7.5, and 10 μg/ml, consistent with one or more exemplary embodiments of the present disclosure. **FIG. 12B** illustrates an evaluation of the toxicity of a commercial CCL21 antigen on MCF7 cancer cells at 24, 48, and 72 hours after incubation using the MTT assay compared to the negative control group at different concentrations of 2.5, 5, 7.5, and 10 μg/ml, consistent with one or more exemplary embodiments of the present disclosure.

### 3.5 Wound healing assay of cytokine based multiepitope protein

The potential activity of the exemplary cytokine-based multi-epitope protein in stimulating the proliferation and migration of cancer cells was investigated using the wound healing assay. FIG. 13 illustrates the effect of the exemplary cytokine-based multi-epitope protein (multi-epitope protein), commercial CCL21 protein, and DMEM medium as a negative control on the migration of MCF7 cancer cells at different times (24, 48, and 72 hours after wound induction), consistent with one or more exemplary embodiments of the present disclosure. Referring to FIG. 13, the photomicrographs of the in-vitro wound assay show that migration and metastasis of cancer cells are pretty evident in control samples and reach about 100% healing after 72 hours. In contrast, the migration rate is low in cells treated with the exemplary cytokine-based multi-epitope protein and commercial protein CCL21 (positive control). In the group treated with the exemplary cytokine-based multi-epitope protein, no noticeable migration of the cells was detected 24 hours after treatment, and 15% and 28% scratch closure rates occurred after 48 and 72 hours, respectively. Furthermore, while the commercial CCL21 showed 90% cancer cell migration after 72 hours, the group treated with the exemplary cytokine-based multi-epitope protein had only 35%, which shows the higher anti-metastatic effect of the exemplary cytokine-based multi-epitope protein than the commercial CCL21. Thus, the MTT assay of and wound healing assay showed that cytokine-based multi-epitope protein has lethality and anti-metastasis effect on MCF7 cancer cells. In other words, the preliminary experiments showed that the exemplary cytokine-based multi-epitope protein could be a suitable choice for experimental cancer research in the future.

**FIG. 13.**
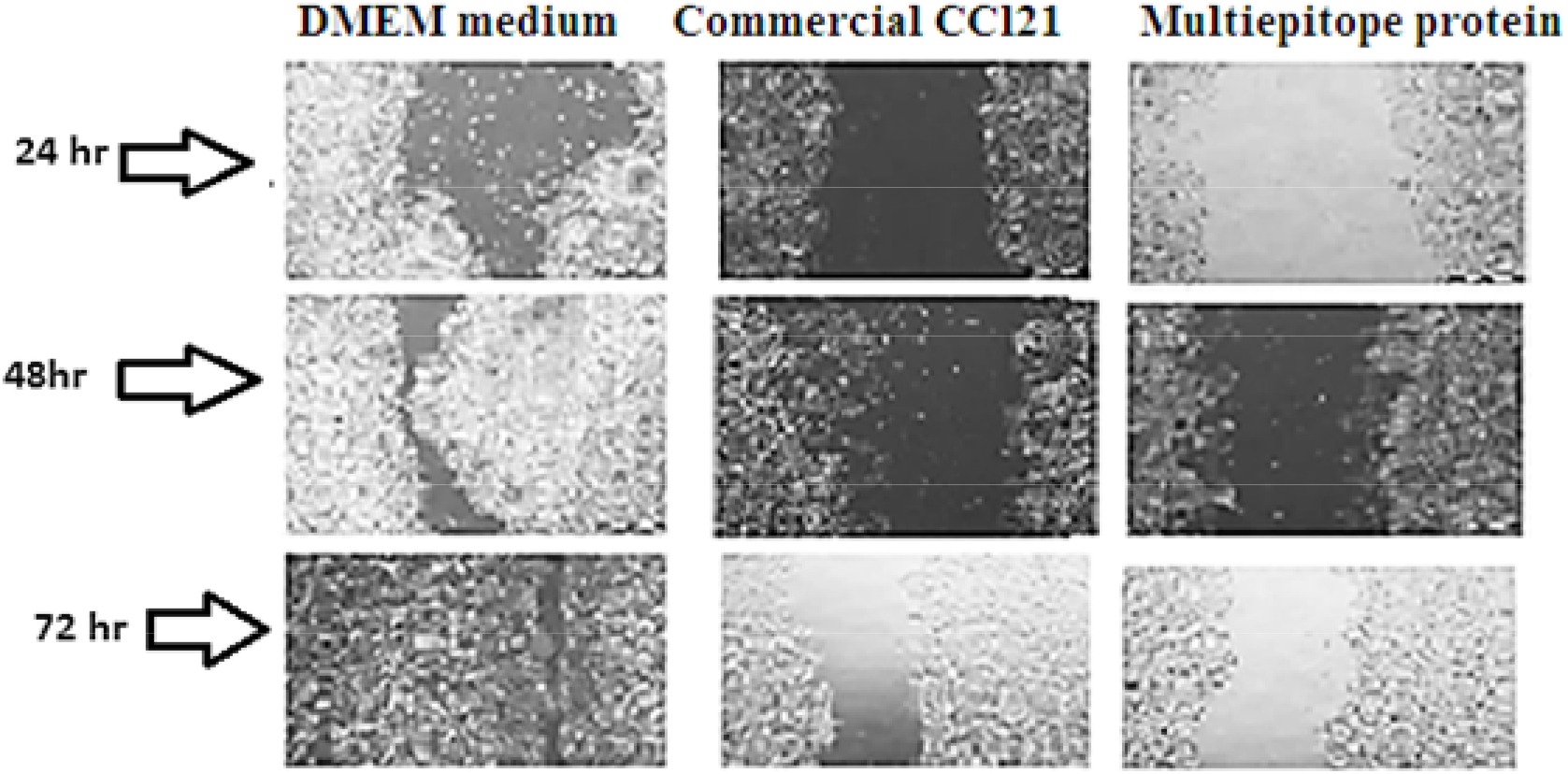
illustrates the effect of the exemplary cytokine-based multi-epitope protein, commercial CCL21 protein, and DMEM medium as a negative control on the migration of MCF7 cancer cells at different times (24, 48, and 72 hours after wound induction), consistent with one or more exemplary embodiments of the present disclosure.

### 3.6 chemotaxis assay of cytokine based multiepitope protein

Chemotaxis assay was used to investigate the effect of CCL21 and CCL19 epitopes of the exemplary cytokine-based multi-epitope protein on T-cell migration since chemokines are cytokines that are involved in leukocyte migration. To observe whether PBMC cells or chemoattractants, including the exemplary cytokine-based multi-epitope protein, attract each other, agarose and Boyden chamber assay was used on CCR7^+^ PBMCs. The fetal bovine serum (FBS) and a commercial CCL21 were used as positive controls. FIG. 14 illustrates a comparison between chemokine (CK) and chemotaxis (CT) properties: A) migration of PBMC cells to the exemplary cytokine-based multi-epitope protein containing CCL21 and CCL19 epitopes (CK), B) migration of PBMC to 10% FBS (CT), consistent with one or more exemplary embodiments of the present disclosure. Referring to FIG. 14, while there is no significant difference between chemotaxis (CT) and chemokine (CK), it can be concluded that the exemplary cytokine-based multi-epitope protein has a chemotaxis effect on the PBMC cells.

**FIG. 14.**
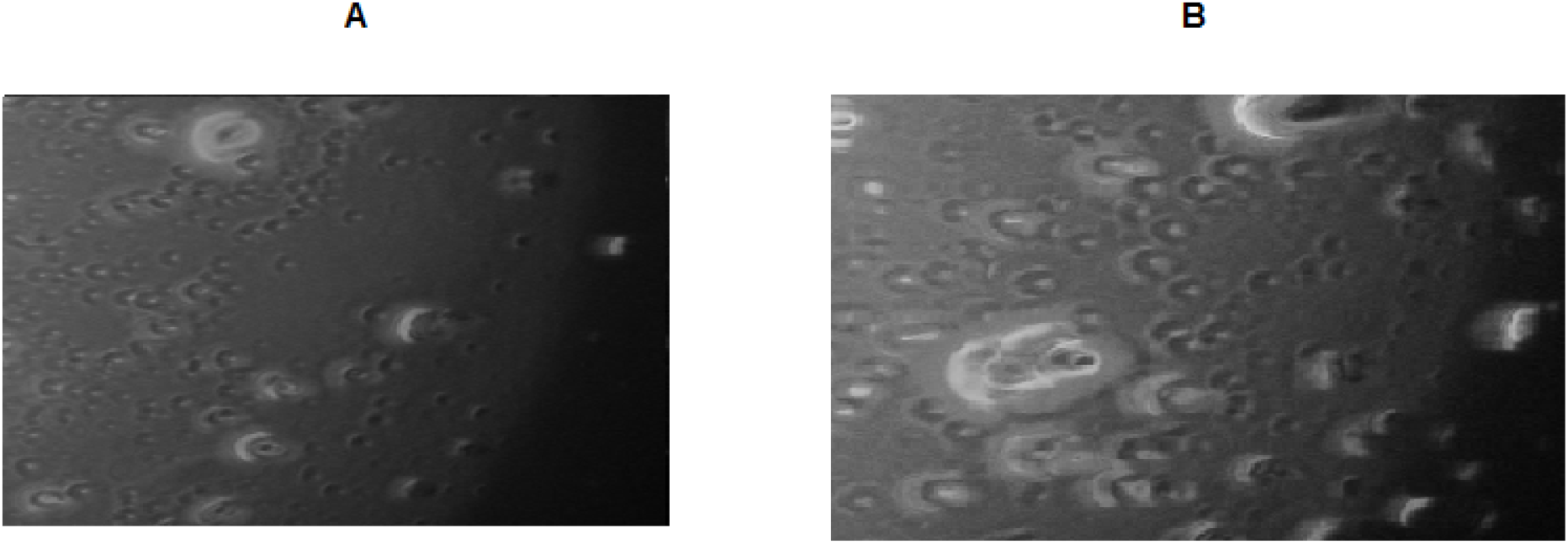
illustrates a comparison between chemokine (CK) and chemotaxis (CT) properties: A) migration of PBMC cells to the exemplary cytokine-based multi-epitope protein containing CCL21 and CCL19 epitopes (CK), B) migration of PBMC to 10% FBS (CT), consistent with one or more exemplary embodiments of the present disclosure.

TABLE 2 represents the chemotaxis rates of different groups treated with the exemplary cytokine-based multi-epitope protein, the commercial CCL21, and FBS. Referring to TABLE 2, the chemotaxis of each substance was assessed by counting the number of cells migrated to each well. As can be seen, the number of monocyte cells moving toward the absorbent agent is higher than the non-absorbent substance, and chemotaxis of the exemplary cytokine-based multi-epitope protein is estimated to be about 91%.

**TABLE 2:** Approximate number of migrating cells and percentage of chemotaxis based on **FIG. 14**

| Adsorbent substance | Cells in the well of adsorbent material | Cells in the well of non-adsorbent material | Chemotaxis rate | Chemotaxis percentage |
| --- | --- | --- | --- | --- |
| Commercial CCL21 | 170 | 17 | 153 | 90 |
| Cytokine-based multi-epitope protein | 138 | 12 | 126 | 91 |
| FBS 10% | 158 | 17 | 141 | 89 |

### 3.7 Evaluation of recombinant protein on animals

By using the CBC test, parameters related to white blood cells such as neutrophils, lymphocytes, basophils, eosinophils, and monocytes can be counted. Therefore, the results shown that the levels of neutrophils, lymphocytes, monocytes, eosinophils and basophils in group B and C, A and D had significant difference (P<0.008). No significant difference was observed in Group A and B and also in group C and group D (P>0.05).

By the above results, we found that intraperitoneal injection of recombinant antigen had an effect on the level of white blood cells in BALB/c mice and acceptable changes were observed in studied groups. Therefore, increase or decrease in the number or type of white blood cells in each of the examined samples (Groups A, …, D) can be a sign of inflammation or cancer, which has caused severe stimulation of the immune system, and the level of white blood cells in tumor mice has increased significantly compared to healthy mice.

## Discussion

In the following detailed description, numerous specific details are presented through examples in order to provide a thorough understanding of the relevant teachings. However, it should be apparent that the present teachings may be practiced without such details. In other instances, well known methods, procedures, components, and/or circuitry have been described at a relatively high-level, without any details, in order to avoid unnecessarily obscuring aspects of the present teachings. The proper scope of the present disclosure may be ascertained from the claims presented below in view of the detailed description and the drawings. In one general aspect, the present disclosure describes an exemplary cytokine-based multi-epitope protein for binding to CC-chemokine receptor type 7 (CCR7)-positive cells(35). In recombinant protein, each of the truncated chemokines may include a DCCL motif, a putative receptor binding cleft, and a putative glycosaminoglycan binding site(35). GM-CSF may be connected to the CCL19 through a helical linker and CCL19 may be connected to the CCL21 through a furine protease-sensitive linker. Also CCL21 may be connected to the truncated IL-1β through a cathepsin-sensitive linker. IL-1β connected to the chemokine secretory signal peptide directly. The chemokine secretory signal peptide may include rat KC chemokine. The CCR7-positive cells may include at least one of CCR7-positive breast cancer cells, CCR7-positive lung cancer cells, monocytes, T lymphocytes, B lymphocytes, natural killer (NK) cells, and dendritic cells (DCs)(38). As a result, the cytokine-based multi-epitope protein may have antitumor properties and may be used for prognosis of tumorigenesis due to higher binding of the CCL21 and CCL19 epitopes of this recombinant protein to the cancer cells with more CCR7 receptors, such as MCF7 cells of breast cancer than the healthy cells. Cytokine-based multi-epitope protein may increase CD8^+^ T cells, leading to better detection and reduction of viral infections progressions(39), such as human immunodeficiency viruses (HIV) and coronavirus disease 2019 (COVID-19) infection. Also it may effectively treat AIDS since the CCL21 epitope of the cytokine-based multi-epitope protein may occupy CCR7 receptors on the surface of CD4^+^ T cells and prevent HIV attachment(38, 40). The IL-Iβ epitope of an exemplary cytokine-based multi-epitope protein may also involve cellular activities, such as neutrophil activation, T- and B-lymphocyte cell production, antibody production, and fibroblast proliferation. Granulocyte-macrophage colony-stimulating factor (GM-CSF) epitope of an exemplary cytokine-based multi-epitope protein as one of the growth factors of white blood cells (WBC) may stimulate stem cells to produce granulocytes and monocytes(22, 41). Additionally, cytokine-based multi-epitope protein may include specific sites for disulfide bonds and linkers to stabilize and activate the cytokine-based multi epitope protein.

